# USVSEG: A robust segmentation of rodents’ ultrasonic vocalization

**DOI:** 10.1101/572743

**Authors:** Ryosuke O. Tachibana, Kouta Kanno, Shota Okabe, Kohta I. Kobayasi, Kazuo Okanoya

**Affiliations:** Department of Life Sciences, Graduate School of Arts & Sciences, The University of Tokyo, Meguro, Tokyo, Japan; Laboratory of Neuroscience, Course of Psychology, Department of Humanities, Faculty of Law, Economics and the Humanities, Kagoshima University; Division of Brain and Neurophysiology, Department of Physiology, Jichi Medical University, Shimotsuke, Tochigi, Japan; Graduate School of Life and Medical Sciences, Doshisha University, Kyoto, Japan

**Keywords:** USV, noise reduction, multitaper method, spectral peak tracking

## Abstract

Rodents’ ultrasonic vocalization (USV) provides useful information to assess their social behaviors. Despite of previous efforts for classifying subcategories of time-frequency patterns of USV syllables to associate with their functional relevances, detection of vocal elements from continuously recorded data have remained to be not well-optimized. We here propose a novel procedure for detecting USV segments in continuous sound data with background noises which were inevitably contaminated during observation of the social behavior. The proposed procedure utilizes a stable version of spectrogram and additional signal processing for better separation of vocal signals by reducing variation of the background noise. Our procedure also provides a precise time tracking of spectral peaks within each syllable. We showed that this procedure can be applied to a variety of USVs obtained from several rodent species. A performance test with an appropriate parameter set showed performance for detecting USV syllables than conventional methods.

## Introduction

Various species in a rodent superfamily *Muroidae* which includes mice, rats, and gerbils, have reported to vocalize ultrasonic sounds in a wide range of frequency up to around 100 kHz [1]. Such ultrasonic vocalization (USV) is expected to be one of useful indicators for describing their social behaviors. Laboratory mice (*Mus musculus domesticus* and *Mus musculus musculus*) have been reported to produce USV for several decades, in particular, as courtship behaviors [2,3]. Their vocalization has known to form a sequential structure [4] which consists of various sound elements, or ‘syllables’. Almost all USV syllables of mice exhibit spectral peaks in 50–90 kHz with having time duration of 10–40 ms, though slight differences in the syllable spectrotemporal pattern were observed among different strains [5]. On the other hand, it has been also well described that laboratory rats (*Rattus norvegicus domesticus*) produce USV syllables which have two predominant categories: one has a relatively higher frequency (around 50 kHz) with a few ten milliseconds in duration, and the other is in the low frequency range (~22 kHz) but much longer duration. These two USV syllables are here named as ‘pleasant’ and ‘distress’ syllables since these are generally considered to be indicators of positive and negative emotional states, respectively [6–9]. This categorization appears to be preserved in different strains of rats, though a slight difference in duration has been reported (Sales, 1979). As another example of the rodent family, vocalizations of Mongolian gerbils (*Meriones unguiculatus*) have also been studied well, in particular, as animal models for audio-vocal system and/or social communication [10–13]. They produce various types of USV syllables ranged up to ~50 kHz with different spectrotemporal patterns [14,15].

In general, rodent’s USV has been estimated to have ecological functions for male-to-female sexual display [2,3,16–19], emotional signal transmission [20–25], and mother-infant interaction [26–29]. Mice USV has been suggested to be distinguished into several subcategories according to their spectrotemporal patterns [30–34], and this pattern could predict mating success [34,35], while the subcategories were not well consistent between those studies. Their USV patterns are innately acquired but not learned behavior [33,36], though sociosexual experience can slightly enhance the vocalization rate [37]. In rats USV, the pleasant (~50 kHz) and distress (~22 kHz) calls have been suggested to have a communicative function since these calls can transmit the emotional states of vocalizer to listener and modify listener’s behavior such as mating [25,38], approaching [39], or defensive behavior [40,41]. Perception of these call also have suggested an effect to modulate listeners’ affective states [20]. Further discrimination of subcategories in the pleasant call has been studied to better know their functional difference in different situations [6–9].

From those characteristics and functions, rodent’s USV is expected to be a good window for sociality and communication studies. Mice USV has been used for studying disorders in social behavior, with a particular focus on autism spectrum disorder [42–45]. Thanks to the recent genetic manipulation technique, USV analysis becomes more important to quantify social behavior of social disorder model mice. On the other hand, studies on rats’ USV have more emphasized the aspect of neural mechanism for the emotional system [6–9], maternal behavior [46–48], or social interaction [49]. USV of other species in the same superfamily of rodents has also been studied as a variety of research models such as parental behaviors, auditory perceptions and vocalization motor controls with gerbils [10,50,51]. Thus, a unified analysis tool for analyzing rodent’s USV is helpful to transfer knowledge obtained across different species.

Previous studies have been proposed analysis toolkits for USV, e.g. MUPET [52] or VoICE [53], and were successful when the recorded sound pieces have enough high signal-to-noise ratio (but mainly focusing on mice’ USV). These analysis tools sometimes suffer from contaminated noises which are produced in normal recording situation such as scratching sounds (short transient noises) and/or stationary background noises produced by fans or air compressors, for example. Such noise greatly deteriorates the segmentation of USV syllables, and also smears acoustical features (e.g. peak frequency) of the segmented syllables, possibly leading to reduction of reliability in classification of vocal categories and quantification of acoustical features of vocalization.

In spite of a variety of behaviors and functions among species, rodent’s USV sound generally tends to exhibit a single salient peak in spectrum with few or none of weak harmonic components (see **Figure 1A** for example). This tendency would be associated with its whistle-like sound production mechanism [54]. From view of sound analysis, this characteristic provides a simple rule for isolating USV sounds from background noises, that is, narrow-band spectral peaks must be vocalized sound while broadband spectral components should be a background. Thus, emphasizing the spectral peaks with flattening the noise floor should improve discrimination of vocalized signal from noise background.

**Figure 1.**
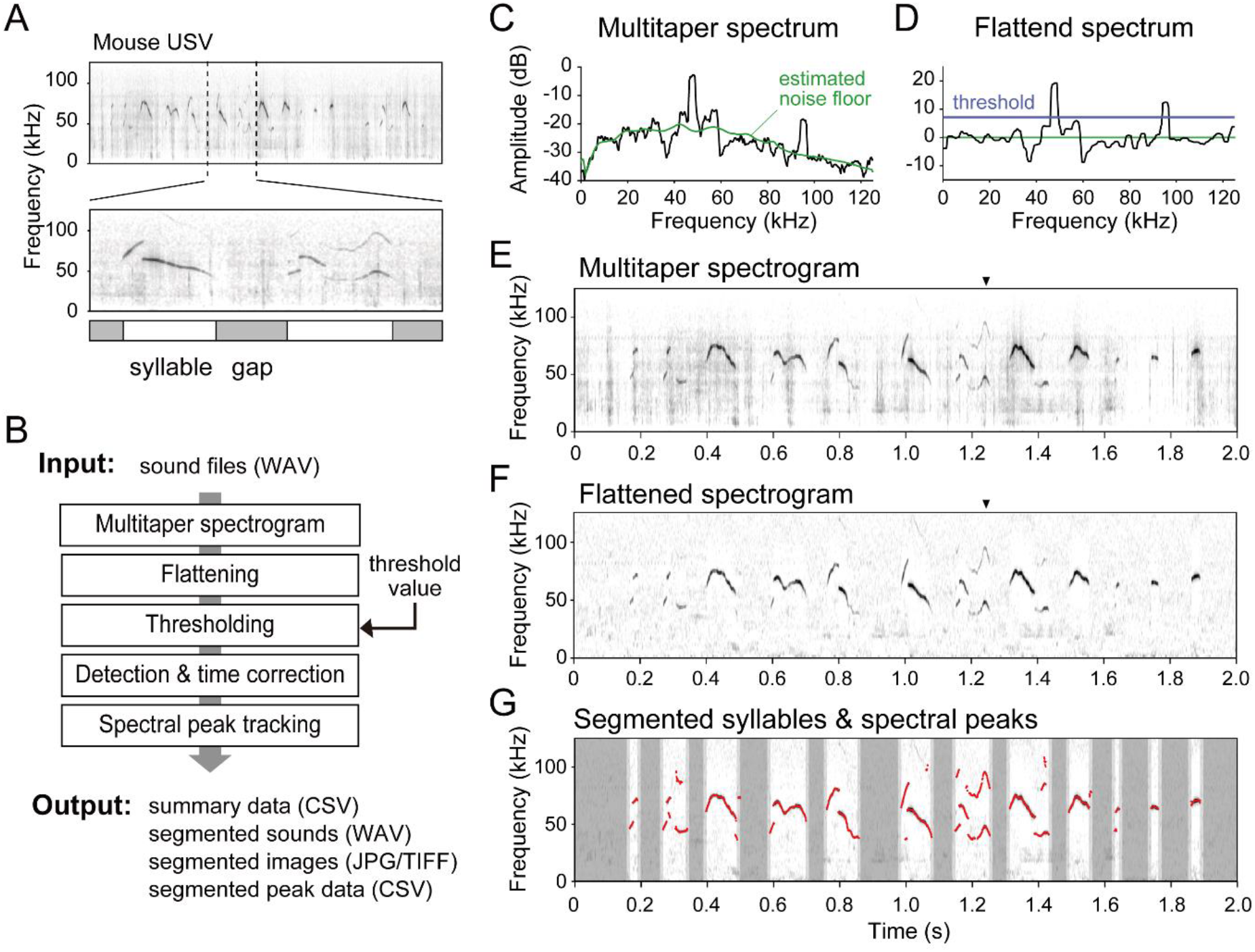
Spectrogram of rodent ultrasonic vocalization (USV) and proposed method for detection of vocal elements in continuously recorded data. **A**. Example spectrogram of mouse vocalization. The brief segment of vocalization (few ten to few hundred milliseconds) is defined as ‘syllable’, and the time interval between two syllables is called as ‘gap’. **B**. Schematic diagram of proposed signal processing procedure. **C-D**. Example of multitaper spectrum and flattened spectrum. **E-F**. Example multitaper (E) and flattened (F) spectrogram obtained from a recording when a male mouse was performing courtship vocalizations to a female mouse. **G**. Processed result of USV data (same as E,F) showing detected syllable periods and spectral peak traces of them (red highlighted dots). Dark gray zone shows non-syllable period and light gray zone indicates a margin inserted before and after syllable period.

We here propose a signal processing procedure for robust detection of USV syllables in recorded sound data by reducing acoustic interference from background noises, and for an additional process to track multiple spectral peaks of the segmented syllables. This procedure consists of five steps (see **Figure 1B**): (1) making a stable spectrogram by the multitaper method which reduces interaction of sidelobes between signal and stochastic background noises, and (2) flattening of the spectrogram by liftering on cepstrum domain which eliminates both pulse-like transient noises and constant background noises. (3) After thresholding, (4) we estimated onset/offset boundaries, and (5) tracked spectral peaks of segmented syllables on the flattened spectrogram. The proposed procedure is implemented in a GUI-based program (“USVSEG”, implemented as MATLAB scripts. https://sites.google.com/view/rtachi/resources), and outputs segmented sound files, image files, and spectral peak feature data after receiving original sound files. These output files can be used for further analyses, for example, clustering, classification, or behavioral assessment by using other toolkits.

The present study demonstrated that the proposed procedure can be applied to a variety of USV syllables produced by a wide range of rodent species successfully, and achieves nearly perfect performance for segmenting syllables in a mouse USV dataset with providing optimized parameter settings. Further, we confirmed that the segmentation performance our procedure is enough higher and more robust against elevated noise than conventional methods.

## Results

### Overview of USV segmentation

Rodents’ ultrasonic vocalization (USV) consist of a series of brief vocal elements, or ‘syllables’ (**Figure 1A**), in a variety of frequency ranges depending on species and situations. For instance, almost all mice strains vocalize in a wide frequency range of 20–100 Hz, while rats’ USV shows a focused frequency around 20–30 kHz when they are distressed. These vocalizations are normally not so prominent and sometime are very unclear in visual inspection of spectrogram because of unavoidable background noises: e.g. fan noises and/or cage noises. Such situation challenges to detection and segmentation of each USV syllable from recorded sound data. Here we assessed a novel procedure consisted of several signal processing methods for segmentation of USV syllables (**Figure 1B**). A smooth spectrogram of recorded sound was obtained using the multitaper method (**Figure 1C,E**) and was flattened by cepstral filtering and median subtraction (**Figure 1D,F**). The flattened spectrogram was binarized with a threshold which was determined in relation to the estimated background noise level. Finally, the signals which exceeded the threshold were used to determine vocalizing period, and integrated to the segmentation (**Figure 1G**). Additionally, our procedure detects spectral peaks every timestep within the segmented syllable periods. In this procedure, users only need to adjust the threshold value for the thresholding process according to signal-to-noise ratio of that recording, while they do not need to adjust other parameters (e.g. maximum and minimum limits of syllable duration and frequency) once they found appropriate values for subjective animals. Note that we provided optimized parameter sets as reference values (see **Table 2**).

### Searching optimal threshold

To assess a relationship between the threshold parameter and segmentation performance, we validated the segmentation performance of our procedure on a mouse USV dataset (**Figure 2**). Actual threshold was defined as multiplication of a weighting factor (or “threshold value”) and the background level, which was quantified as the standard deviation (σ) of an amplitude histogram of the flattened spectrogram (**Figure 2A**; see **Methods** for detail). With low, or high threshold values, the segmentation could miss weak vocalizations, or mistakenly detect noises as syllables, respectively (**Figure 2B**). To find an optimal threshold value for normal recording conditions, we performed a validation test on a dataset including 10 recorded sound files obtained from 7 mice that had onset/offset timing information defined manually by a human expert (**Table 1**). We calculated hit and correct rejection (CR) rates to quantify the consistency of segmentation by the proposed procedure and the manual processing (see **Methods**). We also defined a correct rate as an average of hit and CR rate. The result showed a tendency that the hit rate became lower while the correct rejection went higher with elevation of the threshold value (**Figure 2C**).

**Figure 2.**
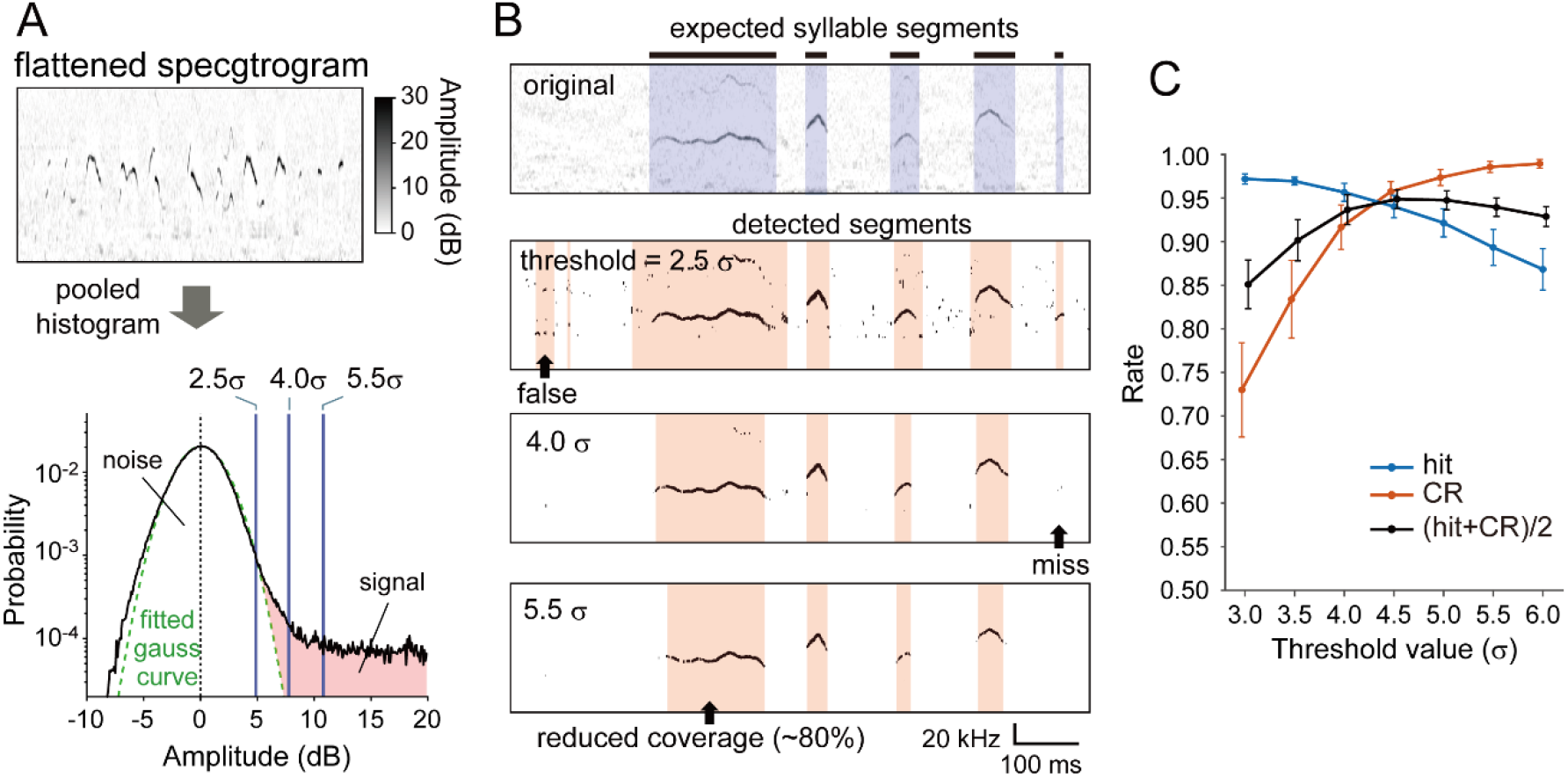
Relationship between thresholding and segmentation performance of mice USV. **A**. Computation scheme of the threshold value. All data points (pixels) of the flattened was pooled and used for making a histogram as a function of amplitude in dB (black line). Background noise distribution was parameterized by a standard deviation (σ) of the gaussian curve (green broken line). The threshold value was defined as a weighting factor of σ (shown as blue vertical lines for example values: 2.5, 4.0 and 5.5). **B**. Example segmentation results for threshold values at 2.5, 4.0 and 5.5 (σ). Uppermost panel shows original flattened spectrogram with segmented periods as syllables by a human expert (blue shaded area). Lower three panels depict thresholded binary images and results of our automatic segmentation for three different threshold values (red shaded). Typically, the amount of false and miss detection increases, and the coverage ratio decreases along with the increase of the threshold value. **C**. Segmentation performance of our procedure as a function of threshold value. Decreasing tendency was observed in the hit rate (blue line), while the correct rejection (CR; red) was increasing with elevation of the threshold value. Average of hit and CR rate (black) was also shown as a balanced index of the performance.

**Table 1.** Rodents USV dataset for performance tests.

| Species/Strain /Condition | Data ID / Subject ID | File duration (s) | # syllables |
| --- | --- | --- | --- |
| Mice (C57BL/6J) | A / Aco59_2 | 113.4 | 333 |
|  | B / Aco59_2 | 115.5 | 85 |
|  | C / Can15-1 | 97.6 | 371 |
|  | D / Can15-1 | 97.0 | 217 |
|  | E / Can16-1 | 135.9 | 394 |
|  | F / Can16-1 | 118.7 | 171 |
|  | G / Can9_2 | 104.7 | 420 |
|  | H / Can9_2 | 205.9 | 306 |
|  | I / Aco65-1 | 98.6 | 383 |
|  | L / Can3_1 | 104.9 | 119 |
| Mice (BALB/c) | BALB128-4 | 60.0 | 168 |
|  | ClnBALB124-4 | 53.0 | 203 |
| Rats (pleasant call) | a14 | 33.2 | 99 |
|  | a18 | 51.6 | 99 |
|  | a42 | 28.7 | 100 |
| Rats (distress call) | a55 | 110.0 | 37 |
|  | a57 | 142.0 | 57 |
|  | a58 | 100.0 | 29 |
|  | a60 | 100.0 | 60 |
| Gerbil | uFMdata | 27.5 | 124 |
|  | uSFMdata | 59.0 | 112 |

### Segmentation performance for various USVs

To demonstrate applicability of the proposed procedure for a wide variety of rodents’ USV, we tested segmentation performance on USV of two strains of laboratory mice (C57BL/6J and BALB/c), two different syllable types in USV of the laboratory rats (PC and DC), and USV syllables of gerbils, respectively. We conducted performance tests for each dataset which included more than 100 syllables with manually detected onset/offset information (**Table 2**). Note that PC and DC in rat vocalization show a remarkable difference in both duration and frequency range even though were produced from identical animals, thus we treated two calls independently and used different parameter sets for them. In the same way as threshold optimization, we calculated hit and CR rate to quantify matching of the automatic segmentations to the manual ones. As results, our procedure showed more than 90% performance in segmentation of various USV syllables (**Figure 3**) with manually tuned parameters (**Table 2**). Slight variability in hit and CR rates was observed across conditions (e.g. the hit rate for gerbil’s USV was lower than that for other conditions), while this should be explained by a difference in the background noise level during recording (as shown in the spectrogram for gerbil’s vocalization containing scratch noises during their vocalizations).

**Figure 3.**
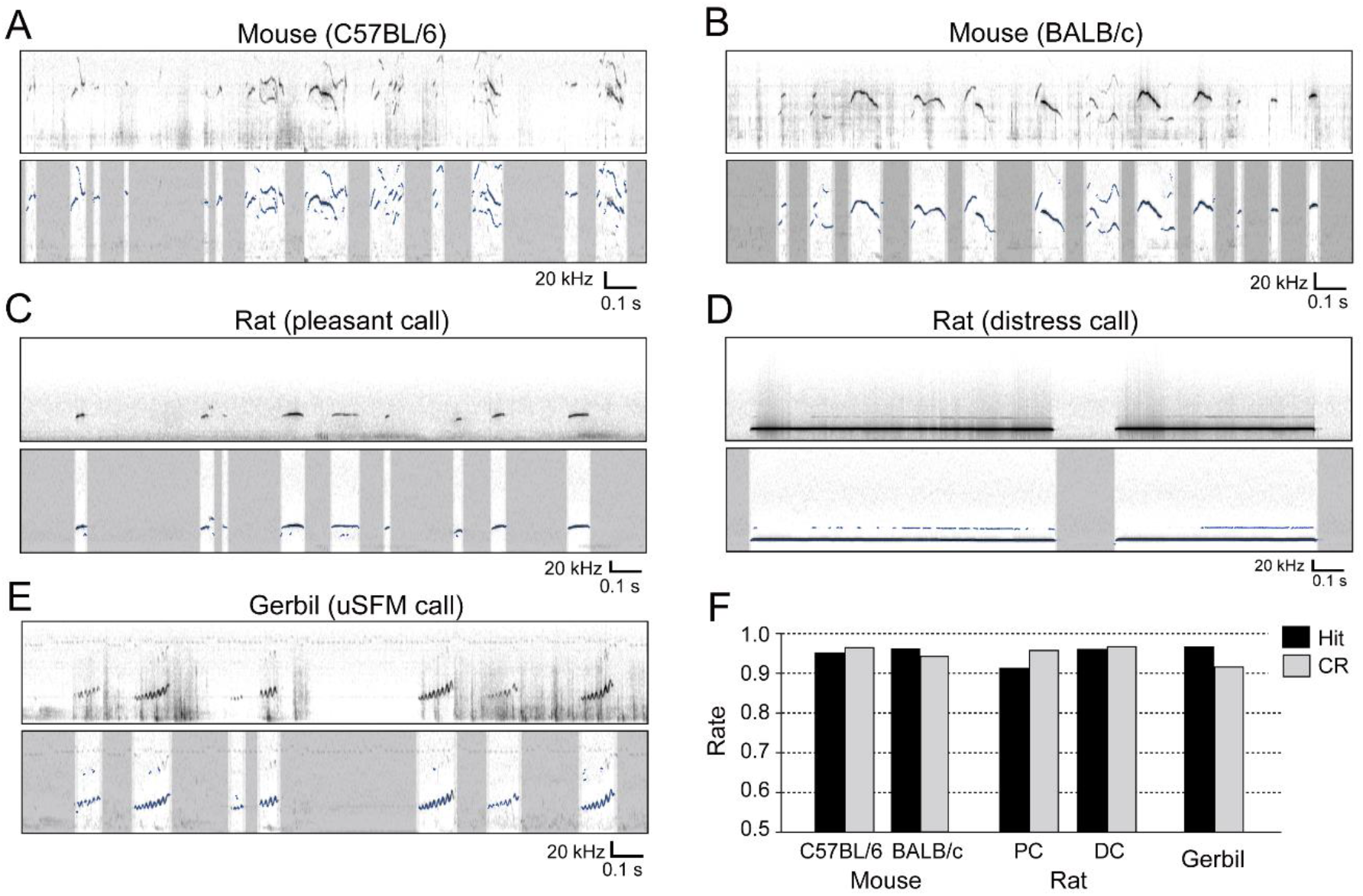
Example results of segmentation and spectral peak tracking on various species in rodents. **A-B**. Mice courtship calls obtained from two different strains: (A) C57BL/6 and (B) BALB/c. **C-D**. Rats calls in context of pleasant (C) and distressed (D) situations. **E**. A representative USV sounds of gerbils, named as upward sinusoidal frequency modulated (uSFM) calls. **F**. Detection performance on all five dataset conditions. All conditions showed more than 90% performance both in the hit rate (black bar) and the correct rejection (CR) rate (gray bar) on the dataset that contains more than 100 syllables (see Method for details).

**Table 2.** Heuristically optimized parameter sets for different species and situations

| Species/Strain /Condition | min. frequency (kHz) | max. frequency (kHz) | min. duration (ms) | max. duration (ms) | min. gap (ms) | threshold value ($\sigma$) |
| --- | --- | --- | --- | --- | --- | --- |
| Mice (C57BL/6J) | 40 | 160 | 3 | 300 | 20 | 4.5 |
| Mice (BALB/c) | 40 | 120 | 3 | 300 | 20 | 4.5 |
| Rats (pleasant call) | 20 | 100 | 3 | 500 | 40 | 3.5 |
| Rats (distress call) | 12 | 40 | 100 | 3000 | 40 | 5.0 |
| Gerbils | 20 | 60 | 5 | 300 | 30 | 4.5 |

### Comparison with conventional methods

We compared our procedure with conventional signal processing methods (**Figure 4**), which were a single-window (or “singletaper”) method for making spectrogram, and the long-term spectral subtraction (“whitening”). The combination of the singletaper and whitening have previously used in other segmentation procedure [55]. According to the study, we used the hanning window as a typical singletaper. Validation tests on the dataset used above were performed with four conditions consisted of combinations of two spectrograms (multitaper vs. singletaper) and two noise reductions (flattening vs whitening). Results demonstrated greater performance with flattening than whitening, and slightly higher in multitaper comparing to singletaper spectrogram (**Figure 4A**). A statistical test showed a significant effect in noise reduction methods (two-way ANOVA; F(1,36) = 31.25, p < 0.001), but the window method effect was not significant (F(1,36) = 0.09, p = 0.770), and no significant interaction between them (F(1,36) = 0.13, p = 0.723). Further, to know robustness of the segmentation methods against noises, we added white noise to the sound dataset at the level of -12, -6, 0, and 6 dB higher than the original sound, and performed the validation test again. We, in particular, compared performance between the multitaper and the singletaper spectrogram with only using flattening for the noise reduction. The result of this test clearly showed that the multitaper method was more robust for the degraded signal-to-noise situations than the singletaper (**Figure 4B**). The statistical test showed significant main effects on both the additive noise level and the window condition with no significant interaction (two-way ANOVA; noise level: F(4,90) = 130.02, p < 0.001; window: F(1,90) = 24.02, p < 0.001; interaction: F(4,90) = 1.29, p = 0.281).

**Figure 4.**
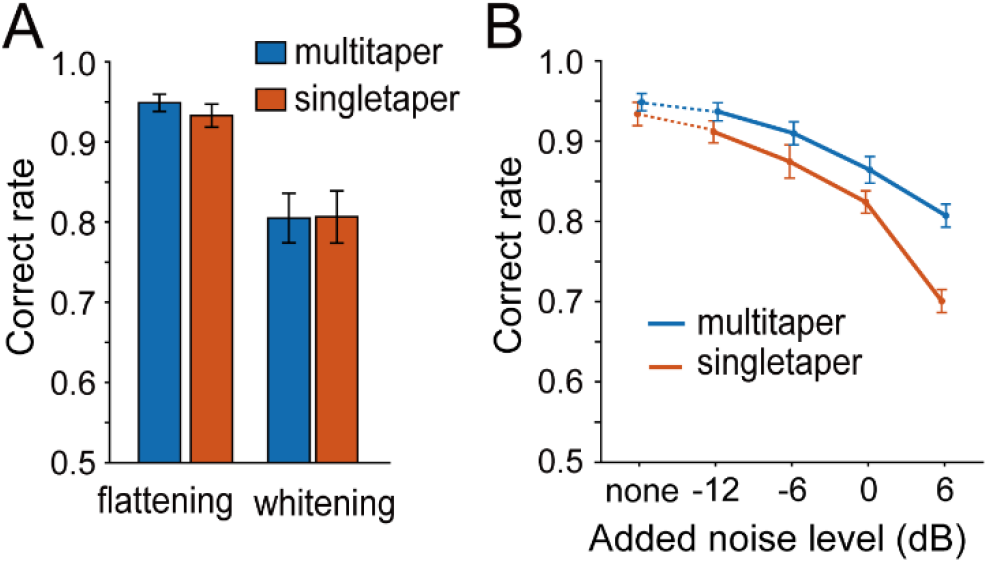
Segmentation performance of various combinations of signal processing methods. **A**. Correct rates of segmentation on flattened or whitened spectrogram produced by multitaper (blue) or single-taper (red) method. We used the hanning window as the single taper condition. **B**. Performance sensitivity to additive noise. Segmentation with multitaper (blue) and single-taper (red) spectrograms against experimentally added background noise. White noise was added to the original data at the level of -12, -6, 0, or 6 dB before processing.

## Discussion

The proposed procedure showed nearly perfect segmentation performance for variable USV syllables of a variety of species and strains in the rodent super family. The procedure was designed to emphasize vocal components in the spectral domain while reducing variability of background noises which usually interfere the segmentation process and inevitably occur during observation of social behavior. This process helped to discriminate vocal signals from the background by thresholding according to the signal-to-noise ratio. Additionally, our procedure also provides a precise tracking of spectral peaks in the vocalization sound. The proposed method set was more robust than the conventional method for syllable detection, in particular, under elevated background noise level. These results demonstrated that this procedure can be generally applied to segment USV of variable rodents.

Our procedure was designed to emphasize distribution difference between vocal signals and background noises, under the assumption that rodent’s USV signal general tendency to have narrow-band sharp spectral peaks. In this process, we employed the multitaper method which uses multiple windows for performing spectral analysis (Thomson 1982), which has been used in vocal sound analyses also for other spices, e.g. songbirds [56]. We also introduced the spectral flattening process in which the broadband spectrum in each timestep was flattened by cepstral filtering. As we demonstrated in the performance comparison tests, a combination of the multitaper windowing with the flattening showed better performance than the conventional method with a single-taper windowing and long-term spectral subtraction that has been used in a mouse USV analysis [55], in particular, under the degraded signal-to-noise conditions. Note that our experimental results did not focus on applicability for lower-frequency (i.e. <20 kHz) harmonic-rich, or harsh noise-like vocalizations since those sounds are outside of our presupposition for the processing algorithm.

We here employed redundant way to represent the spectral feature of USV, and exporting 1up to 3rd candidates of spectral peaks for every timestep. This provides additional information about harmonics, and also an appropriate way to treating “jumps” which are sudden change in the spectral peak tracks [4]. Researchers have tried to distinguish syllables into several subcategories according to their spectrotemporal features to know sequential patterns of them [31,34,35,57]. The present study will lead to a future possibility to have better categorization of USV syllable subtypes.

## Methods

### Proposed procedure

Our procedure consisted of five steps: multitaper spectrogram, flattening, thresholding, segmentation, and spectral peak tracking. In particular, the multitaper method and the flattening were core processes for suppressing variability of background noise as described below in detail.

#### Multitaper spectrogram

We used the multitaper method [58] for obtaining spectrogram to improve signal saliency against background noise distribution. Multiple time windows (or tapers) were designed as a set of 6 series of the discrete prolate spheroidal sequence with setting the time half bandwidth parameter to 3 [59]. The length of these windows was set to 512 samples (~2 ms for 250 kHz sampling rate). In each time step, the original waveform was multiplied with all the six windows and transformed to the frequency domain. Derived six spectrograms were averaged into one to obtain a stable spectrotemporal representation. This multitaper method reduces background noises comparing to usual single-taper spectrogram, while widening bandwidth of signal spectral peaks.

#### Flattening

To emphasize spectral peaks for detectability of vocalization events, we reduced variability of background noises by flattening the spectrogram. This flattening consists of two processes. First, transient broadband (or impulse-like) noises were reduced by the liftering in every time step, in which a gradual fluctuation in frequency domain, or spectral envelope was filtered out by replacing the first three cepstral coefficients to zero. This process can emphasize spectral peaks of rodent’s ultrasonic vocalization since they have few or none of harmonics and are very narrow-band. Then, we calculated a grand median spectrum that had median values of each frequency channel, and subtracted it from the liftered spectrogram.

#### Thresholding and detection

After the flattening, we binarized the flattened spectrogram image at a threshold which was determined based on estimated background noise level. The threshold was calculated as multiplication of a weighting factor (or “threshold value”) and the standard deviation (σ) of a background distribution (**Figure 3A**). The σ value was estimated from a pooled amplitude histogram of the flattened spectrogram as described in the previous study for determining onsets and offsets of birdsong syllables [60]. The threshold value was normally chosen from 3.5–5.5 and could be manually adjusted depending on the background noise level. After binarization, we counted the maximum number of successive pixels along with frequency axis whose amplitude exceeded the threshold in each time frame, and regarded the time frame as to include vocalized sounds when the counted maximum number was 5 or more (corresponded to a half bandwidth of the multitaper window).

#### Timing correction

A pair of detected elements split by a silent period (or gap) with duration less than a predefined lower limit (“gap min”) was integrated to omit unwanted segmentation within syllables. We usually set this lower limit for gap around 3–30 ms according to animal species or strains. Then, the sound elements with duration of more than a lower limit (“dur min”) were judged as syllables. If the duration of element was exceeded the upper limit (“dur max”), then the element was excluded. These two parameters (dur min and max) were differently determined for different species, strains, or situations. The heuristically optimized values of these parameters were suggested in a table (**Table 2**) as reference.

#### Spectral peak tracking

We also implemented an algorithm for tracking multiple spectral peaks as additional analysis after the segmentation. Although the focus of our study is temporal segmentation of syllables, we briefly explain this algorithm as follows. First, we calculated saliency of spectral peaks by convolving a second-order differential spectrum of the multitaper window itself into the flattened spectrum along with frequency axis. This process emphasis steepness of spectral peaks in each time frame. Then, the strongest four local maxima were detected in the spectral saliency as candidates. We grouped the four peak candidates to form a continuous spectral object according to their time-frequency continuity (within 5% frequency change per a time frame). If a length of the grouped object was less than 10 time points, then spectral peak data in the object were excluded from candidates for vocalized sound. At the final step, the algorithm outputs maximum three peaks in each time step.

### Dataset

For testing the segmentation performance of our procedure, we prepared datasets consisted of recorded sounds and manually detected onset/offset timings of syllables for three species in the rodents superfamily (**Table 1**). The manual segmentations for different species were performed by different human experts. These experts segmented sound materials by visual inspection of spectrogram, independently of any automatic segmentation system. They were not informed any result of our procedure of that material beforehand. We finally collected segmented data for more than 100 syllables in total from each condition (species, strains, or contexts) as below. For all species/strain/context conditions, ultrasonic sounds were recorded using a commercial condenser microphone and an A/D converter (Ultra-SoundGate, Avisoft Bioacoustics, Berlin, Germany; SpectoLibellus2D, Katou Acoustics Consultant Office, Kanagawa, Japan). The whole dataset is available from a website (https://sites.google.com/view/rtachi/resources).

#### Mice

We obtained 10 recording sessions of courtship vocalizations from 6 mice (*Mus musculus;* C57BL/6J, adult males), under the same condition and recording environment as described in our previous work (Kanno and Kikusui 2018). In brief, the microphone was set 16 cm above the floor with a sampling rate of 400 kHz. Latency to the first call was measured after introducing a same strain adult female into the cage and then ultrasound recording was performed for an additional minute. The data recorded during the first minute after the first ultrasound call were analyzed for the number of calls. For all the recording tests, the bedding and cages for the males were exchanged one week before the recording tests, and these home-cage conditions were maintained until the tests were completed. These sound data files were recorded for other works (in preparation) originally, and shared on mouseTube (https://mousetube.pasteur.fr) and Koseisouhatsu Data Sharing Platform (http://data-share.koseisouhatsu.jp). Note that we chose 10 files from the shared data with excluding two files (“J” and “K”) which does not contain enough number of syllables for the present study. For obtaining data from an animal of the other strain (BALB/c), we used the same procedure with C57BL/6J, but with a sampling rate of 250 kHz.

#### Rats

The pleasant call (PC) or distress call (DC) was recorded from an adult female rat for each (*Rattus norvegicus domesticus*; LEW/CrlCrli, Charles River Laboratories Japan). For the recording of PC, the animal was given massage-like stroking stimuli on the experimenter’s lap with hand for around 5 minutes. To elicit DC vocalization, another animal was transferred to a wire-topped experimental cage and habituated to the cage for 5 minutes, then, received air puff stimuli (0.3 MPa) with an inter-stimulus interval of 2 s to the nape from a distance of approximately 5–10 cm. Immediately after 30-times air puss stimuli, ultrasonic vocalizations were recorded for 5 min. These vocalizations were detected by a microphone placed at a distance of approximately 15–20 cm from the target animal. The detected sound was digitally recorded at a sampling rate of 384 kHz.

#### Gerbils

Vocalizations of the Mongolian gerbil (*Meriones unguiculatus*) were recorded via the microphone positioned 35 cm above an animal cage, and the cage was in the center of a soundproof room. The sound was digitized at a sampling rate of 250 kHz. These sound data were originally obtained from our previous study [14]. We here targeted only calls whose fundamental frequencies were in the ultrasonic range (20 kHz or more), i.e. upward FMs and upward sinusoidal FMs, which were often observed under conditions of what appeared to be mating and a non-conflict context [14].

### Performance test

#### Quantification of segmentation performance

We calculated two indices for segmentation performance named as hit and correct rejection (CR) rate. The hit rate corresponds to a probability of true-positive detection, which shows how many true syllables (defined by the human expert) were detected in our procedure. When the duration of detected syllable covered more than 90% of the true syllable period (defined by human expert), the detection was counted as the hit. We defined the hit rate as a ratio of the number of hits against the number of true syllables. The CR rate indicates a probability of true-negative detection, meaning how many detected syllables were not false positive. We first calculated a ratio of false detection (less than 1% overlap with true syllable periods) against the total number of detections. Then, we defined the CR rate as 1 – false detection rate.

#### Threshold optimization

The segmentation would miss weak vocal sounds when the threshold is too high, or mistakenly detect noises as syllables with the higher threshold. To find an optimal threshold value for normal recording conditions, we assessed segmentation performance on a dataset for mice USVs with changing the threshold value. For this test, we used 10 files of C57BL/6J mice from the dataset. We varied the threshold value from 3.0 to 6.0 with 0.5 step. After obtaining of the hit and CR rate for each threshold value, we additionally calculated average of hit and CR rate: (hit + CR)/2, as a balanced performance index that was expected to be the maximum when the threshold value was optimal.

#### Comparison with conventional method

To know to what extent our procedure improved the detection performance from conventional methods, we compared the performance of four conditions in which two processing steps were swapped with conventional ones. As conventional methods, we employed a normal windowing method (“singletaper” condition) using the hanning window as a replacement of the multitaper method for making spectrogram, and the long-term spectral subtraction (“whitening” condition) for replacing with the flattening process. These two methods have used as standard processing methods for such signal detection algorithms [55]. Here we swapped either or both of two windowing and two noise reduction methods between ours and conventional ones for making four conditions: multitaper+flattening, multitaper+whitening, singletaper+flattening, and singletaper+whitening. Then, performance of them was tested on the dataset of mice USV that was the same one used for the threshold optimization test. Note that the threshold for bandwidth in the detecting process after the binarization was adjusted to 3 for the singletaper method (it was 5 for the multitaper) to be corresponded to its half bandwidth.

#### Noise addition

As further analysis for assessing the robustness against noise, we carried out the performance test again on the multitaper-flattening and singletaper+flattening conditions with adding white noise to the original sound data at the level of -12, -6, 0, and 6 dB referring to the root-mean-square of the original sound amplitude. We have not tested the whitening method here since the method showed clearly lower performance than the flattening in the performance test without adding noise.

## Acknowledgement

The authors thank Yui Matsumoto, Koh Takeuchi, Kazuki Iijma, and Yumi Saito for helpful comments and feedbacks for developing the software, and critical discussions on an earlier version of the manuscript.

## Supporting information

This work was supported by MEXT/JSPS KAKENHI Grant No. 16H06528 to KK, 17H05956 to SO, 18H05089 to KIK, and 17H06380 to KO.

## Ethical information

All procedure for vocalization recording were approved by the Ethics Committee of Azabu University (#130226-04) for mice, and the Animal Experiment Committee of Jichi Medical University (#17163-02) for rats.

## Author contribution

ROT designed the study, developed the proposed method, performance validation, and wrote the manuscript. KK collected mice data, manual segmentation of them, feedback for developing the proposed method, and supported in the preparation of the manuscript. SO collected rats data, manual segmentation of them, and feedbacked to the manuscript. KIK collected gerbils data, manually segmented them, and feedbacked to the manuscript. KO supervised entire study, and feedback to the manuscript. All authors approved the final version of the manuscript, and agree to be accountable for all aspects of the work in ensuring that questions related to the accuracy or integrity of any part of the work are appropriately investigated and resolved.

